# How to identify the best index case in families with hereditary breast and ovarian cancer

**DOI:** 10.1101/527168

**Authors:** Margot J. Wyrwoll, Lea Fuchs, Daniel E.J. Waschk

**Affiliations:** Institute of Human Genetics, University Hospital Münster, Vesaliusweg 12-14, 48149 Münster (GER)

**Author notes:** **Correspondence to:** Daniel E.J. Waschk (MD PhD), Institute of Human Genetics, University Hospital Münster, Vesaliusweg 12-14, 48149 Münster (GER). **Funding:** The German Consortium for Hereditary Breast and Ovarian Cancer is funded by the German Cancer Aid (110837). **Abbreviations:** BC: breast cancer; OC: ovarian cancer; HBOC: hereditary breast and ovarian cancer; GC-HBOC: German Consortium for Hereditary Breast and Ovarian Cancer; CI: confidence interval. **Definitions:** *Mutation detection rate* refers to the detection of expected mutations according to pretest risk calculations; *Mutation prevalence* refers to the proportion of patients with a mutation of the whole cohort.

**Keywords:** Hereditary, Breast Cancer, Ovarian Cancer, Best Index Case, Patient Selection, Mutation Detection Rate

## Abstract

To date, a disease-causing mutation can be found in approximately 15-30% of families with hereditary breast and ovarian cancer and still more than half of the cases remain unsolved. Usually it is intended to perform genetic analyses in the family member with the most severe phenotype, which, however, is not always possible. Moreover, no standard criteria have been established to define the person who is most suitable for genetic testing within a family: *the best index case*. This study now establishes clinical selection criteria to identify the best index case in families with hereditary breast and ovarian cancer and analyses the impact on genetic testing. 130 patients who presented at our department from 2016 to 2018 were divided into two groups. In group A, genetic analyses were performed in the best index case (N = 98). In group B, at least one family member had a more severe phenotype compared to the person who was tested (N = 32). The mutation detection rate was significantly higher for group A compared to group B (64.3% vs. 32.0%, p = 0.034), even though there was no significant difference of calculated mutation carrier risks between these groups. Furthermore, the mutation detection rate in group A was notably higher compared to the results of previous studies. We conclude that the mutation detection rate in families with hereditary breast and ovarian cancer can be improved by identifying the best index case for genetic testing according to the clinical selection criteria reported here and suggest that these can be used as a guideline for genetic counseling.

## Introduction

Approximately 12.8% of all women in Germany are affected by breast cancer (BC) throughout their lives [1]. Thus, BC represents the most common type of cancer in women. The lifetime risk for male breast cancer is estimated at about 0.1% [1], the risk for ovarian cancer (OC) at 1.4% [2]. Familial clustering suggestive of hereditary breast and ovarian cancer (HBOC) is seen in 30% of cases [1]. Mutations in the high-penetrance susceptibility genes *BRCA1* (OMIM 113705) and *BRCA2* (OMIM 600185) are well known causes for HBOC [3]. Lately, associations of BC with mutations in other genes such as *CHEK2* (OMIM 604373), *PALB2* (OMIM 610355) and *CDH1* (OMIM 192090), among others, have been described additionally [4, 5].

In the last years, standard genetic analyses for HBOC within a clinical setting covered ten genes in Germany [6]. Identification of a disease-causing mutation can be of major importance for the treatment and surveillance of affected persons as well as for healthy relatives [1-3, 6]. Currently there are 21 centers of the German Consortium for Hereditary Breast and Ovarian Cancer (GC-HBOC) which attend affected patients and their families [7]. Genetic counseling is performed for all cases and includes risk stratification with the software BRCAPRO. This software implements a statistical model based on Mendelian genetics and Bayes’ theorem for calculating an individual’s probability of carrying a mutation on the basis of the individual’s cancer status and the reported family history [8, 9]. According to most studies, a disease-causing mutation can be found in 15-30% of families with HBOC and it is believed that more than half of the cases still remain unsolved [10-12]. Divergent inclusion criteria and *a priori* mutation carrier risks are the main reasons for the different mutation prevalences, which makes it difficult to compare the results. Not only the inclusion criteria for genetic testing but also the selection criteria for the patient who is most suitable for genetic testing within a family, henceforward *the best index case/patient*, varies among the different studies. Moreover, if the best index case of a family is not available, genetic analyses are frequently performed in other family members. There usually exists more than one affected person with BC and/or OC in families suggestive of HBOC. Not all of them must necessarily carry the disease-causing mutation which would be expected to be present in the family. Occasionally, family members can also be affected by a cancer which developed independently of the family burden. Therefore, correct selection of the best index case for genetic testing is crucial in order to detect a disease-causing mutation. Usually it is intended to perform genetic analyses in the family member with the ‘most severe’ phenotype depending on the number and type of cancers (either BC and/or OC) and the age of onset [10]. However, no standard criteria have been established to clearly define and rank the severity of these cases so far, leaving the decision at the discretion of the respective genetic counselor. In some families identification of the index patient with the most severe phenotype can be straightforward. However, in many other families this is not the case. As an example, the question whether a woman with a single BC at age 41 (younger age of onset) or a woman with OC at age 49 (older age of onset but less frequent type of cancer) should be chosen for genetic analyses within a family needs to be addressed by ranking the severity of these cases. Furthermore, it is of interest if a standardized approach for the selection of the best index cases can improve the results of genetic analyses in terms of mutation detection rates.

The present study therefore establishes clinical selection criteria applicable to most families suggestive of HBOC to identify the best index case and analyses the impact on genetic testing. To achieve this goal 130 patients who presented at our clinic from 2016 to 2018 were divided into two groups. In group A, genetic analyses were performed in the best index patient according to the criteria in table 1. In group B, at least one other family member was considered a better index patient. The results of genetic analyses were then compared for both groups with the average mutation risk probabilities according to BRCAPRO calculation.

**Table 1.**
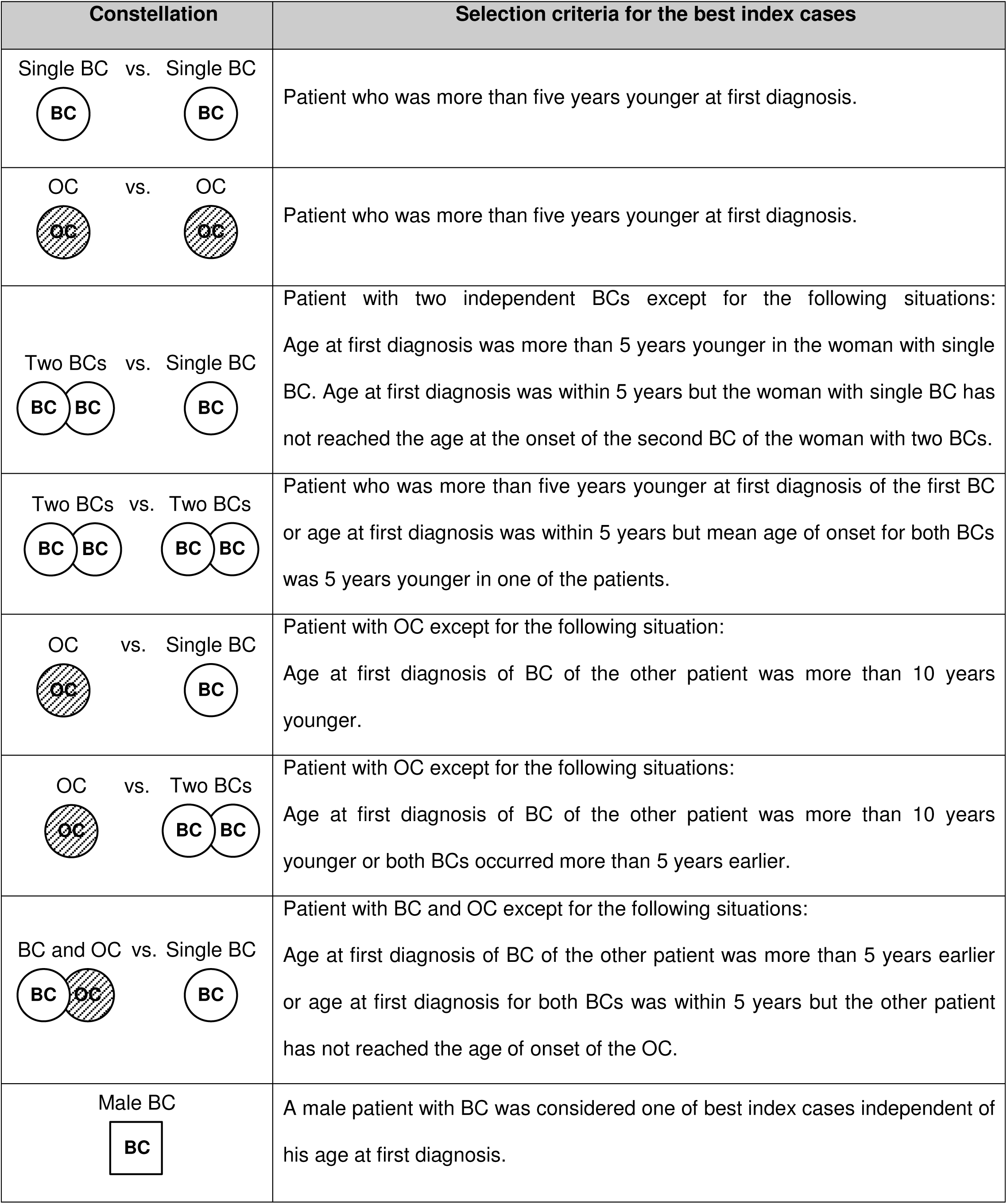
Clinical selection criteria that were applied in this study to identify the best index case for genetic analyses in families with hereditary breast and ovarian cancer. BC: Breast cancer; OC: Ovarian cancer

| Constellation | Selection criteria for the best index cases |
| --- | --- |
| Single BC vs. Single BC | Patient who was more than five years younger at first diagnosis. |
| OC vs. OC | Patient who was more than five years younger at first diagnosis. |
| Two BCs vs. Single BC | Patient with two independent BCs except for the following situations:<br>Age at first diagnosis was more than 5 years younger in the woman with single BC. Age at first diagnosis was within 5 years but the woman with single BC has not reached the age at the onset of the second BC of the woman with two BCs. |
| Two BCs vs. Two BCs | Patient who was more than five years younger at first diagnosis of the first BC or age at first diagnosis was within 5 years but mean age of onset for both BCs was 5 years younger in one of the patients. |
| OC vs. Single BC | Patient with OC except for the following situation:<br>Age at first diagnosis of BC of the other patient was more than 10 years younger. |
| OC vs. Two BCs | Patient with OC except for the following situations:<br>Age at first diagnosis of BC of the other patient was more than 10 years younger or both BCs occurred more than 5 years earlier. |
| BC and OC vs. Single BC | Patient with BC and OC except for the following situations:<br>Age at first diagnosis of BC of the other patient was more than 5 years earlier or age at first diagnosis for both BCs was within 5 years but the other patient has not reached the age of onset of the OC. |
| Male BC | A male patient with BC was considered one of best index cases independent of his age at first diagnosis. |

## Materials and Methods

### Patients

The study includes 130 Caucasian patients and their families from Germany who presented at the center for HBOC at the University Hospital Münster for genetic counseling for the first time from 2016 to 2018. Informed consent was obtained from all participants and all cases fulfilled the requirements of the GC-HBOC to offer genetic testing [7, 10]. The software Cyrillic (v2.1) was used for pedigree drawing and pretest mutation risk calculation with BRCAPRO based on the reported family history. Families were selected for the study by the criteria shown in supplementary table 1 to ensure reliable pretest risk calculation.

### Selection of the best index cases

The 130 families were divided into two groups depending on whether genetic testing was performed in one of the best index cases of a family (Group A, N = 98) or whether there was at least one affected family member that was more suitable but not available for genetic testing (Group B, N = 32). The clinical selection criteria shown in table 1 were applied in this study in order to determine if there was a better index case within a family. Accordingly, the woman with OC at age 49 from the example in the introduction would be the best index patient since the age of onset was less than 10 years later compared to the woman with BC.

### Genetic analyses

Sequence analyses were performed by next-generation sequencing on an Illumina MiSeq/NextSeq system for the coding exons and adjacent intronic regions of the genes *BRCA1, BRCA2, RAD51C, CHEK2, PALB2, ATM, RAD51D, NBN, CDH1* and *TP53* according to the specified genes of the GC-HBOC [6]. Coverage was at least 100 and Phred-Score was at least 30 for all nucleotides. All detected mutations were confirmed by Sanger sequencing. In four cases of group A and one case of group B sequence analyses were limited to the genes *BRCA1, BRCA2, RAD51C, CHEK2* and *PALB2* after a disease-causing mutation was identified in one of these genes. For the genes *BRCA1* and *BRCA2* multiplex ligation-dependent probe amplification (MLPA; MRC-Holland) was additionally performed in all cases to detect larger alterations such as deletions and duplications of whole exons. Mutations classified as pathogenic or likely pathogenic according to the guidelines of the International Agency for Research on Cancer [13] and the American College of Medical Genetics [14] were considered as disease-causing.

### Statistics

The software IBM SPSS statistics (v25) was used for statistical analyses. Comparison of the average pretest mutation carrier probability for the index cases of groups A and B was performed with Mann–Whitney *U* test. The difference of correct results and false negative results (missed mutations) according to pretest risk calculations between groups A and B was analyzed with chi-square test. Values for p < 0.05 were considered statistically significant.

## Results

Average mutation carrier probability was 33.3% (95% CI: 26.8-39.8) for group A and 39.1% (95% CI: 27.4-50.8) for group B (Table 2). Mutation carrier probability was 34.7% (95% CI: 29.1-40.4) for the whole cohort. The difference of average carrier probabilities between groups A and B was statistically not significant (p = 0.386). According to pretest calculations, the expected number of patients with a disease-causing mutation was 32.7 for group A and 12.5 for group B. A disease-causing mutation was found in 21 patients of group A and in 4 patients of group B. The mutation detection rate was significantly higher (p = 0.034) in group A (64.3 %) compared to group B (32.0%) (Table 2, Figure 1). The distribution of the mutations per gene is summarized in supplementary table 2. One patient of group A carried a disease-causing mutation both in *CHEK2* and *CDH1.*

**Table 2.** Results of pretest risk calculations and genetic analyses. Group A: Genetic testing was performed in the best index case. Group B: At least one affected family member was more suitable for genetic testing.

|  | Group A | Group B | Total |
| --- | --- | --- | --- |
| **Number of Patients** | 98 | 32 | 130 |
| **Average mutation**<br>**carrier risk** | 33.3%<br>(95% CI: 26.8-39.8) | 39.1%<br>(95% CI: 27.4-50.8) | 34.7%<br>(95% CI: 29.1-40.4) |
| **Expected number of**<br>**patients with a mutation** | 32.7 | 12.5 | 45.2 |
| **Number of patients with**<br>**a detected mutation** | 21 | 4 | 25 |
| **Mutation**<br>**detection rate** | 64.3% | 32.0% | 55.3% |

**Figure 1.**
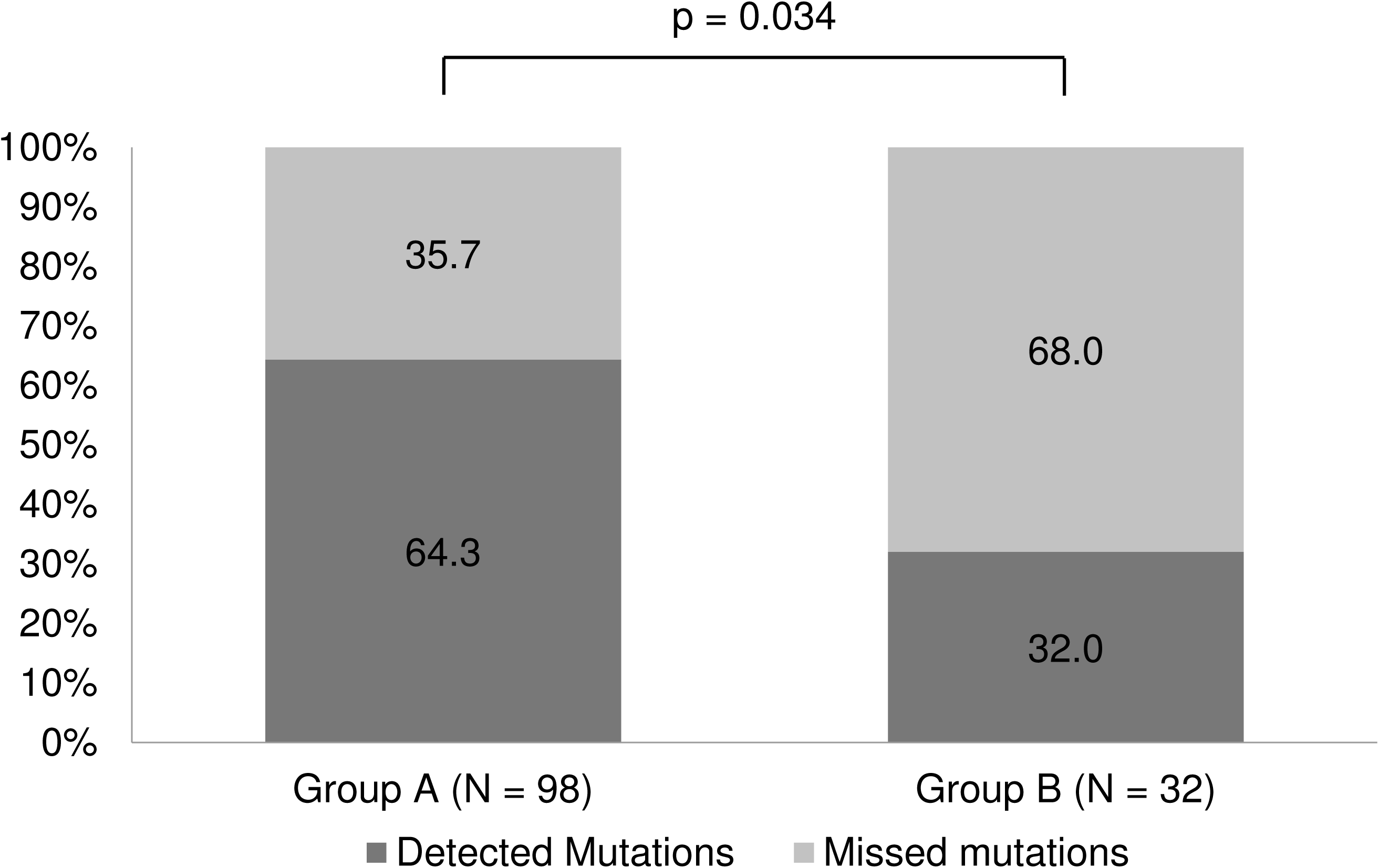
Mutation detection rates for groups A and B. Group A: Genetic testing was performed in the best index case. Group B: At least one affected family member was more suitable for genetic testing.

## Discussion

A recently published review summarized that approximately 25% of HBOC cases can be solved by genetic analyses when taking into account the high- and moderate risk genes which are covered by the analyses of the present study [15]. Compared to this number, the mutation detection rate of the present study is notably higher (64.3%) for the group with the best index cases, suggesting that the outcome of genetic analyses can be improved notably by using the selection criteria reported here. As shown in the example from the introduction, the best index patient must not necessarily be the person with the youngest age of onset. Instead, it can also be a patient with the less frequent type of cancer. Without standardized criteria, the selection of the best index basically depends on the choice of the genetic counselor and therefore can vary significantly.

Of course, in some families the best index patient may not be available or declines genetic analyses. In other cases direct testing of a different person can be justified if the cancer treatment depends on an early result. Nevertheless, identification and selection of the best index case for genetic analyses based on the criteria reported here should be emphasized to optimize the testing results. If genetic analyses were already performed in another family member without detecting a disease-causing mutation, additional genetic testing for the best index case should be offered in order to not miss a mutation in the family. It should also be investigated if blood or tissue samples are still available if the best index patient is already deceased.

Patients were selected precisely for this study according to the enrollment criteria found in supplementary table 1 to ensure reliable calculations of mutation carrier risks. The downside of this approach is the limitation of the sample size of the cohort, which, however, does not affect statistical significance of the results. We therefore suggest to perform a larger multicenter trial to confirm our findings and further validate our selection criteria for clinical practice. Although statistically not significant, the average mutation carrier risk was slightly higher for group B compared to group A (39.1% vs. 33.3%, p = 0.386). This may seem paradox since the patients of this group are not the best index cases of their respective families. However, both the average mutation carrier risk and the probability that there can be found a better index case raise simultaneously with the number of persons affected by BC/OC within a family. In this context we additionally calculated the average mutation carrier risk for the best index patients of group B who were not available for genetic testing. We found no noteworthy difference between them and the tested patients of group B (41.9% vs. 39.1%, p = 0.672), which shows the limitation of using the mutation carrier risk alone for finding the best index patient. In group B 68.0% of the expected mutations were missed compared to only 35.7% in group A (Figure 1), suggesting that a reasonable number of women with BC/OC do not carry the disease-causing mutation which would be expected to be present in the family. An explanation for this could be that some family members are affected by a cancer which developed independently of the family burden. This option seems to be probable especially when taking into account the high incidence of sporadic BC in the population. Mutations in yet undiscovered susceptibility genes or imprecise information about affected family members are further explanations that still not all families with HBOC seem to be solved by genetic analyses today.

Also, statistical models as used in BRCAPRO produce only an estimation of mutation carrier risks. Nevertheless, pretest risk calculation for the tested cohorts is essential to be able to compare the outcome of genetic analyses between different studies. BRCAPRO is currently the only software in clinical use by the GC-HBOC since it has the highest diagnostic accuracy [16] and alternative programs such as BOADICEA have not been approved for clinical practice yet. At the time BRCAPRO was designed, *BRCA1/2* were the only susceptibility genes known for HBOC. When considering mutations in *BRCA1/2* only, the mutation detection rate is also about double as high for group A in this study. Independent of this, the creators stated that other susceptibility genes would be also covered by BRCAPRO if disease penetrances were comparable to those of *BRCA1/2* [9], which can be assumed at least for *TP53* [15, 17], *PALB2* [1, 5] and *CDH1* [17].

In summary, this study establishes clinical selection criteria which can be applied easily to most families suggestive of HBOC in order to identify the best index case for genetic testing. Mutation detection rates are significantly higher when genetic analyses are performed in the best index case compared to other affected family members. We suggest that the selection criteria reported here can be used as a guideline for genetic counseling of HBOC.

## Acknowledgements

We thank all patients and their families for participating in this study.

## Author contribution

### Margot Wyrwoll

Selection of patients, data collection, drafting of manuscript.

### Lea Fuchs

Preparation of pedigree charts for genetic counseling.

### Daniel E.J. Waschk

Conception and design, genetic counseling, BRCAPRO risk calculation, selection of patients, data collection, statistical analysis, drafting of manuscript.

### All authors

Data analysis and interpretation, critical revision and final approval of manuscript.

## Conflict of Interest

The authors declare that they have no conflict of interest.

---

**Supplementary Table 1.** Inclusion and exclusion criteria for study enrollment of patients toensure reliable pretest risk calculation with BRCAPRO. BC: Breast cancer; OC: Ovarian cancer; HBOC: Hereditary breast and ovarian cancer

| Inclusion criteria |
| --- |
| • Family history was complete for at least three generations including ages of onset of BC/OC cases and current ages of unaffected family members.<br>• Genetic analyses were performed in a family member affected by BC and/or OC as stated in the methods section.<br>• Pathology reports on the subtype of cancer were available at least for the tested person. |
| Exclusion criteria |
| • Incomplete knowledge about ages of onset and types of cancer in affected persons as well as number and current ages of unaffected relatives within three generations of the family.<br>• Pathology reports were not available for the tested person.<br>• Pathology reports were conflicting when trying to distinguish a relapse from a *de novo* cancer in cases with metachronous breast cancer.<br>• Consanguinity was reported for the ancestors of the tested person.<br>• A disease-causing mutation for HBOC was already known in the family at first presentation.<br>• Other conditions leading to an increased risk of breast cancer were present in the tested person such as radiation of the thorax in the treatment of Hodgkin's Lymphoma or genetic conditions like Neurofibromatosis type 1. |

**Supplementary Table 2.** Number of detected mutations for each gene and for thecategories ‘*BRCA1/2*’ and ‘Other genes’. †One patient from group A carried a disease-causing mutation both in *CHEK2* and *CDH1.*

| Category | Number of Mutations |  |  |
| --- | --- | --- | --- |
|  | Group A | Group B | Total |
| *BRCA1/2* | 14 | 3 | 17 |
| Other genes | 8 | 1 | 9 |
| Gene | Group A | Group B | Total |
| *BRCA1* | 9 | 3 | 12 |
| *BRCA2* | 5 | 0 | 5 |
| *CHEK2*<sup>†</sup> | 5 | 1 | 6 |
| *RAD51C* | 0 | 0 | 0 |
| *PALB2* | 2 | 0 | 2 |
| *RAD51D* | 0 | 0 | 0 |
| *ATM* | 0 | 0 | 0 |
| *CDH1*<sup>†</sup> | 1 | 0 | 1 |
| *NBN* | 0 | 0 | 0 |
| *TP53* | 0 | 0 | 0 |

## References

1 S3-Guideline on Diagnostics, Therapy and Follow-up of Breast Cancer (2018) German Guideline Programme in Oncology. Version 4.1. https://www.leitlinienprogramm-onkologie.de/leitlinien/mammakarzinom/ Accessed 7 January 2019

2 S3-Guideline on Diagnostics, Therapy and Follow-up of Malignant Ovarian Tumours (2018) German Guideline Programme in Oncology. Version 3.01. https://leitlinienprogramm-onkologie.de/Ovarialkarzinom.61.0.html Accessed 7 January 2019

3 Petrucelli N, Daly MB, Pal T (2016) BRCA1-and BRCA2-Associated Hereditary Breast and Ovarian Cancer. GeneReviews® [Internet]. Seattle (WA): University of Washington, Seattle; 1993–2018. 1998 Sep 4 [updated 2016 Dec 15]. https://www.ncbi.nlm.nih.gov/books/NBK1247/ Accessed 7 January 2019

4 Bernstein JL, Teraoka SN, John EM, Andrulis IL, Knight JA, Lapinski R, Olson ER, Wolitzer AL, Seminara D, Whittemore AS, Concannon P (2006) The CHEK2*1100delC allelic variant and risk of breast cancer: screening results from the Breast Cancer Family Registry. Cancer Epidemiol Biomarkers Prev 15:348–352. https://doi.org/10.1158/1055-9965.EPI-05-0557

5 Antoniou AC, Casadei S, Heikkinen T et al (2014) Breast-cancer risk in families with mutations in PALB2. N Engl J Med 371:497–506. https://doi.org/10.1056/NEJMoa1400382

6 Schmutzler R (2017) Konsensusempfehlung des Deutschen Konsortiums Familiärer Brust- und Eierstockkrebs zum Umgang mit Ergebnissen der Multigenanalyse. Geburtshilfe und Frauenheilkunde 77:733–739. https://doi.org/10.1055/s-0043-108531

7 German Consortium for Hereditary Breast and Ovarian Cancer (2019) https://www.konsortium-familiaerer-brustkrebs.de Accessed 10 April 2019

8 Parmigiani G, Berry D, Aguilar O (1998) Determining carrier probabilities for breast cancer-susceptibility genes BRCA1 and BRCA2. Am J Hum Genet 62:145–158.

9 Berry DA, Iversen ES Jr, Gudbjartsson DF, Hiller EH, Garber JE, Peshkin BN, Lerman C, Watson P, Lynch HT, Hilsenbeck SG, Rubinstein WS, Hughes KS, Parmigiani G (2002) BRCAPRO validation, sensitivity of genetic testing of BRCA1/BRCA2, and prevalence of other breast cancer susceptibility genes. J Clin Oncol 20:2701–2712.

10 Kast K, Rhiem K, Wappenschmidt B et al (2016). Prevalence of BRCA1/2 germline mutations in 21 401 families with breast and ovarian cancer. J Med Genet 53:465–471. https://doi.org/10.1136/jmedgenet-2015-103672

11 Machackova E, Foretova L, Lukesova M, Vasickova P, Navratilova M, Coene I, Pavlu H, Kosinova V, Kuklova J, Claes K (2008) Spectrum and characterisation of BRCA1 and BRCA2 deleterious mutations in high-risk Czech patients with breast and/or ovarian cancer. BMC Cancer 8:140. https://doi.org/10.1186/1471-2407-8-140

12 Frank TS, Deffenbaugh AM, Reid JE, Hulick M, Ward BE, Lingenfelter B, Gumpper KL, Scholl T, Tavtigian SV, Pruss DR, Critchfield GC (2002) Clinical characteristics of individuals with germline mutations in BRCA1 and BRCA 2: analysis of 10,000 individuals. J Clin Oncol 20:1480–1490.

13 Plon SE, Eccles DM, Easton D, Foulkes WD, Genuardi M, Greenblatt MS, Hogervorst FB, Hoogerbrugge N, Spurdle AB, Tavtigian SV (2008) Sequence variant classification and reporting: recommendations for improving the interpretation of cancer susceptibility genetic test results. Hum Mutat 29:1282–1291. https://doi.org/10.1002/humu.20880

14 Richards S, Aziz N, Bale S, Bick D, Das S, Gastier-Foster J, Grody WW, Hegde M, Lyon E, Spector E, Voelkerding K, Rehm HL (2015) Standards and guidelines for the interpretation of sequence variants. a joint consensus recommendation of the American College of Medical Genetics and Genomics and the Association for Molecular Pathology. Genet Med 17:405–424. https://doi.org/10.1038/gim.2015.30

15 Wendt C, Margolin S (2019) Identifying breast cancer susceptibility genes - a review of the genetic background in familial breast cancer. Acta Oncol 3:1–12. https://doi.org/10.1080/0284186X.2018.1529428

16 Fischer C, Kuchenbäcker K, Engel C et al (2013) Evaluating the performance of the breast cancer genetic risk models BOADICEA, IBIS, BRCAPRO and Claus for predicting BRCA1/2 mutation carrier probabilities: a study based on 7352 families from the German Hereditary Breast and Ovarian Cancer Consortium. J Med Genet 50:360–367. https://doi.org/10.1136/jmedgenet-2012-101415

17 Mahdavi M, Nassiri M, Kooshyar MM, Vakili-Azghandi M, Avan A, Sandry R, Pillai S, Lam AK, Gopalan V (2018) Hereditary breast cancer; Genetic penetrance and current status with BRCA. J Cell Physiol 234: 5741–5750. https://doi.org/10.1002/jcp.27464

